# Childhood socioeconomic status moderates genetic predisposition for peak smoking

**DOI:** 10.1101/336834

**Authors:** Laura Bierut, Pietro Biroli, Titus J. Galama, Kevin Thom

## Abstract

Smoking is the leading cause of preventable disease and death in the U.S., and it is strongly influenced both by genetic predisposition and childhood socioeconomic status (SES). Using genetic variants exhibiting credible and robust associations with smoking, we construct polygenic risk scores (PGS) and evaluate whether childhood SES mediates genetic risk in determining peak-cigarette consumption in adulthood. We find a substantial protective effect of childhood SES for those genetically at risk of smoking: adult smokers who grew up in high-SES households tend to smoke roughly the same amount of cigarettes per day at peak (∼ 23 for low and ∼ 25 for high genetic risk individuals, or about 8%more), while individuals from low-SES backgrounds tend to smoke substantially more if genetically at risk (∼ 25 for low and ∼ 32 for high genetic risk individuals, or about 28% more).

## 1 Introduction

Smoking is the leading cause of preventable disease and death in the U.S. (Mokdad et al., 2004). Each year tobacco use kills nearly 440,000 Americans – who die up to 15 years earlier than nonsmokers – and costs more than $193 billion in annual health-related economic losses (Mokdad et al., 2004; DHHS, 2010; Kaplan et al., 1995). Smoking is more prevalent among low socioeconomic status (SES) groups (Cutler and Lleras-Muney, 2010) and a significant source of the substantial disparities in health between them (Chaloupka and Warner, 2000; Contoyannis et al., 2004; Mackenbach et al., 2004; Khang et al., 2009; Cutler et al., 2011). Such disparities are formed early in life and become more pronounced as individuals age (Case et al., 2002, 2005; Currie and Stabile, 2003; Shonkoff et al., 2009). Indeed, an extensive literature has shown the importance of early-life circumstances in explaining adult health outcomes (Currie and Rossin-Slater, 2015; Caspi et al., 2016; Doyle et al., 2009; Institute of Medicine, 2015; Osler et al., 2003; Almond and Currie, 2011; Condliffe and Link, 2008; Fernald et al., 2012; Gluckman and Hanson, 2006b,a; Bateson and Gluckman, 2011).

Genetic makeup matters too. Twin studies (comparing the correlation in traits between identical and fraternal twins) suggest that some 35% to 86% of the variance in heaviness of smoking is related to genetic differences between individuals, with the remainder attributed to environmental influences (Kaprio et al., 1981; Swan et al., 1990; Hettema et al., 1999; Koopmans et al., 1999; Lessov et al., 2004; Broms et al., 2006; Boardman et al., 2010).

While genes may predispose individuals to certain unhealthy behaviors and to certain health conditions, the influence of genetic factors depends substantially on environmental exposures, so called gene-by-environment (G×E) interplay (Haldane, 1946; Plomin et al., 1977; Rutter,2006; McAllister et al., 2017). Theory suggests SES disparities emerge by an interaction of cumulative disadvantage and genes, pointing at the importance of the early-life environment (Kuh and Shlomo, 2004; Caspi, 2004; Meaney and Szyf, 2005; Meaney, 2010; Cole, 2009; Rutter, 2012; Mitchell et al., 2013; Reiss et al., 2013). Therefore, G×E interplay may be cumulative in nature, and stronger for early-vs. late-life environments.

Thus far, however, studies have focused largely on G×E interplay between contemporaneous environments and smoking (Boardman et al., 2010, 2011; Fletcher, 2012; Domingue et al., 2015;Meyers et al., 2013; Schmitz and Conley, 2016; Treur et al., 2017). For example, one recent study (Meyers et al., 2013) suggests that an individual ‘s *current* social environment in adulthood moderates genetic vulnerability to smoking.^1^

Here, we test the hypothesis that a protective socioeconomic environment during childhood moderates the effect of genetic risk for smoking in adulthood. Two recent developments make our analysis possible. The study of G×E interplay has traditionally been hampered by a lack of credible gene-behavior associations and by a lack of data sets with both genetic and precisely measured data of the socioeconomic environment. Studies attempting to discover genetic main effects or G×E interaction were severely underpowered to detect true associations (Benjamin et al., 2012; Duncan and Keller, 2011; Hewitt, 2012; McGue, 2013). Recent advances in molecular genetics, however, have established robust associations between specific genetic variants and smoking behavior (Liu et al., 2010; Thorgeirsson et al., 2008, 2010; The Tobacco and Genetics Consortium et al., 2010). These studies used significantly larger samples, applied stringent standards for statistical significance, and demonstrated out-of-sample replication to robustly identify genetic associations.

A second critical development has been the recent collection of genetic samples in large, representative data sets containing extensive measures of socioeconomic environment, health, and health behavior, at different points in the life cycle. Such variables are typically not found in the medical samples used for gene discovery, and social science data sets have lacked genetic information. We here use rich information from one such study, the Health and Retirement Survey (HRS) (Sonnega et al., 2014).

We provide empirical evidence that high childhood SES substantially reduces genetic risk for peak life-cycle levels of smoking. This supports the notion that growing up in a nurturing socioeconomic environment can offset the genetic risk of heaviness of smoking.

## 2 Results

Single nucleotide polymorphisms (SNPs) differ across individuals, thereby providing a measure of genetic variation. Following strict statistical procedure for multiple hypotheses testing (*p* < 5 × 10^−8^) and controlling for population stratification, previous genome-wide association studies (GWAS) have identified genome-wide significant relationships between specific SNPs and smoking quantity (Liu et al., 2010; Thorgeirsson et al., 2010; The Tobacco and Genetics Consortium et al., 2010). We focus here on a measure of heaviness of smoking that is highly correlated with nicotine use disorder, the number of cigarettes smoked per day at peak consumption (max CPD). To maximize statistical power, we aggregate SNPs (associated with the CPD phenotype in an independent sample) with *P*-values < 10^−4^ into a standardized polygenic risk score (PGS) for max CPD (std. dev. = 1, mean = 0), using the combined GWAS coefficients in Table 1 of (Thorgeirsson et al., 2010) as weights. We work with an HRS sample of 3,280 ever-smokers, born between 1920 and 1960, with non-missing genetic and demographic data, and who are genetically of European descent. This follows accepted practices of restricting the genetic diversity to match as closely as possible the genetic profile of the discovery sample. In the HRS, max CPD is a retrospective item, defined only for ever-smokers. We find a positive and highly statistically significant (*p* = 1.57 × 10^-3^) relationship between the PGS and max CPD, with each additional unit of the PGS associated with 0.90 extra cigarettes per day.

**Table 1:**
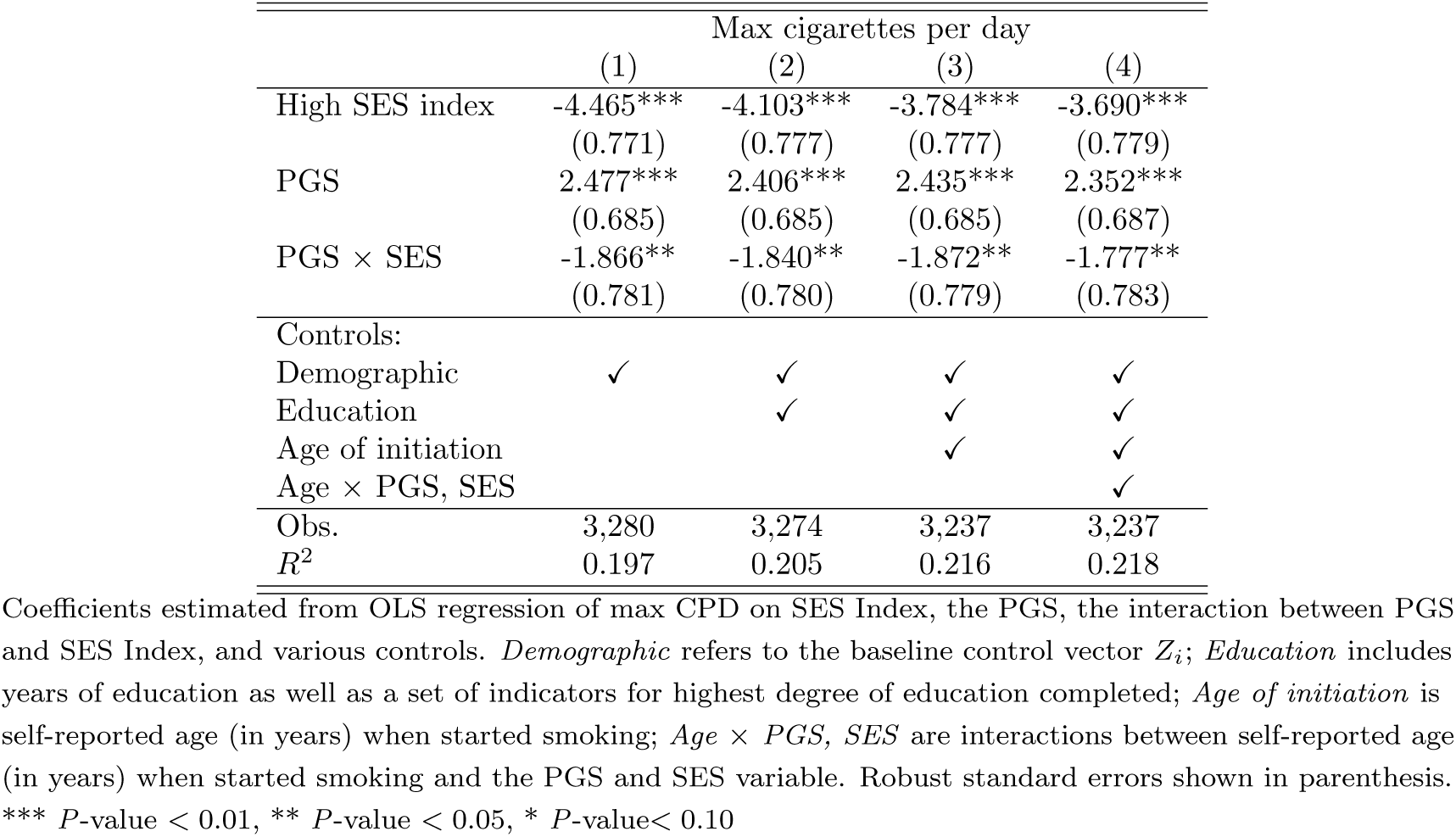
Interplay between PGS and high SES index in max CPD

From recall questions, we generated three distinct dichotomous variables (taking values{0, 1 }) capturing higher SES from birth till age 16. The three childhood SES binaries indicate: (1) Family Well Off: above-average or average family financial position in childhood, (2)Never Move or Help: the family never had to move or ask relatives for help for financial reasons during childhood, and (3) Father Employed: the individual never endured an extended period with an unemployed or absent father (see Supporting Information [SI] Section 4.2 for detail). As a fourth dichotomous childhood SES variable, we created an index that sums over the three binary SES measures (taking values {0, 1, 2, 3 }) and divided the sample into low SES (values{0, 1 }) and high SES (values {2, 3 }).

We regressed max CPD on indicators for PGS and high SES, along with their interaction,of the following form:

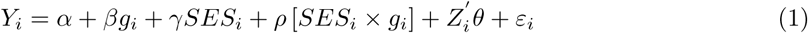

where *Y*_*i*_ is max CPD for individual *i*; *g*_*i*_ is the polygenic risk score; *SES*_*i*_ is one of the four dichotomous measures of household SES during childhood, and *Z*_*i*_ is a vector of individual characteristics. The coefficient *ρ* measures the extent to which SES moderates the association between the PGS and smoking.

### Analysis

Figure (1) shows the unconditional relationship in the raw data between max CPD and genetic risk (PGS, expressed in std. dev. from mean zero) for individuals from low- and high-SES backgrounds (based on the index measure of SES). Differences in the slope of the relationship between the two SES groups is evidence supporting the existence of a G×E interaction. For a high childhood SES background (dashed line) there is little difference in max CPD between high and low genetic risk, while for a low childhood SES background (solid line) max CPD rises significantly with increasing genetic risk. Those from a high SES background smoked around 23 (low genetic predisposition, PGS = -2) or 25 (high genetic predisposition, PGS = +2) cigarettes per day at peak, while those from a low SES background smoked around 25 (low PGS)or 32 (high PGS), respectively. Thus, the genetic gradient is small (∼ 8%) for those from high childhood SES backgrounds and large (∼ 28%) for those from low childhood SES backgrounds. These differences are statistically significant, as the non-overlapping 99% confidence intervals illustrate: there is strong evidence of interaction between genetic predisposition and childhood SES.

**Figure 1:**
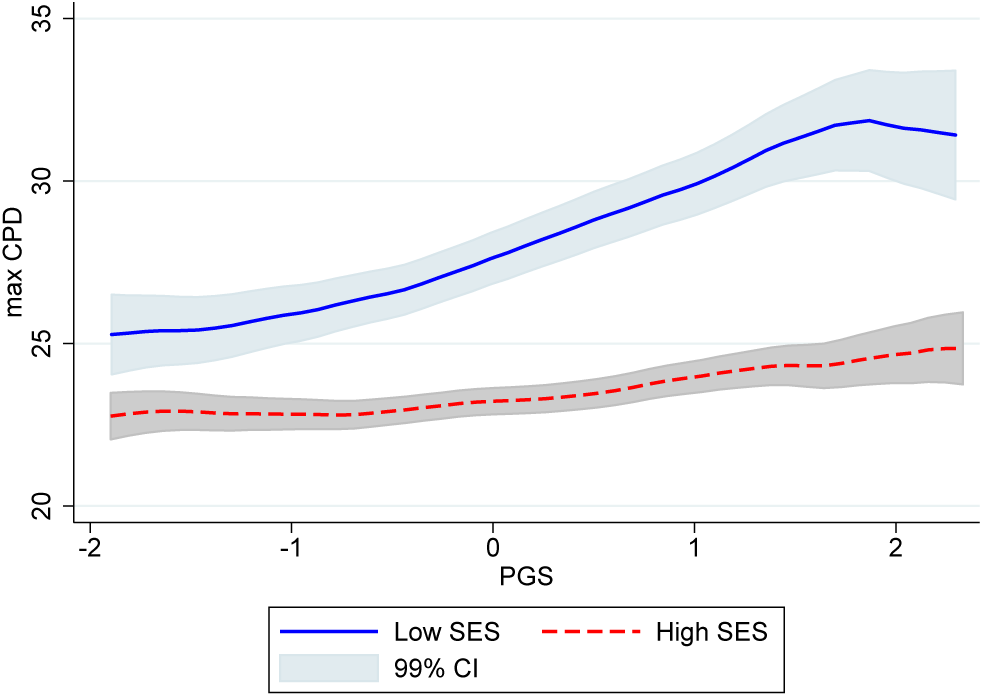
Max cigarettes per day (max CPD) as a function of the polygenic risk score (PGS) for low (solid blue) and high (dashed red) childhood SES Index. The shaded area indicates the 99 percent confidence interval. In constructing the Figure, we used kernel-weighted local polynomial smoothing. All estimations were performed in STATA using the command *lpoly*. To avoid outliers, the sample was trimmed below the 1^*st*^ and above the 99^*th*^ percentile of max CPD and the PGS. Note that the PGS distribution is slightly skewed to the right. As a result, the x-axis is not symmetric around zero.

Next, we investigate whether these results are robust to controlling for individual characteristics such as cohort, gender, region of birth, education, and population stratification. Throughout our analyses, our baseline control vector *Z*_*i*_ includes the first ten principal components of the full matrix of SNP data to adjust for population stratification (Price et al., 2006; Rietveld et al., 2014), a male indicator, a set of indicators for birth year, a set of indicators for region of birth (New England, Middle Atlantic, East North Central, West North Central, South Atlantic, East South Central, West South Central, Mountain, and Pacific), interactions between the male indicator and birth year indicators, between the male indicator and region of birth indicators, and between the birth year indicators and region of birth indicators.

Table 1 reports the OLS estimations of equation (1) using different sets of controls. We find that both high childhood SES and low genetic predisposition reduce peak smoking. Further, the coefficient on the interaction term, *ρ*, is consistently negative, and statistically significant. To interpret these results, consider our baseline specification in column 1. The coefficient estimates suggest that, on average, an individual from a low-SES background with low genetic risk (PGS1 std. dev. below mean) smokes 4.95 (2 × 2.477) fewer cigarettes per day at peak than an individual from a low-SES background with high genetic risk (PGS 1 std. dev. above mean). However, this genetic gradient almost vanishes for those from high-SES backgrounds: a highgenetic-risk individual smokes on average only 1.22 more cigarettes per day at peak than a low genetic risk individual.^2^

Now contrast the results with education. A college graduate smokes, on average, 4.30 (s.e. 0.93) fewer max CPD than a high-school dropout.^3^ The interaction between low SES and high genetic risk (2 × 1.866 = 3.732 max CPD) is comparable in size to the difference in max CPD between a college graduate and a high-school dropout. Education is generally considered of great importance in the social sciences, associated with significant differences in health, health behaviors, income, and longevity (Hauser and Kitagawa, 1973; Christenson and Johnson, 1995; Elo and Preston, 1996; Rogers et al., 2000, 2010; Lleras-Muney, 2005; Conti et al., 2010; Cutler et al., 2011; Clark and Royer, 2013; Meghir et al., 2018). Thus, the observed G×E interaction is not only statistically significant but also economically meaningful.

As shown by columns (2), (3) and (4), these results are robust to the inclusion of potential confounding factors, such as educational attainment and age of smoking initiation. The results are also robust to including full interactions between the PGS, SES, and the set of controls *Z*_*i*_ (as suggested by Keller, 2014; see SI Tables 6 and 7). The difference in coefficients across columns is always very small and never statistically significant, even when including very predictive controls (Oster, 2016). For instance, age of smoking initiation is significantly correlated with SES and a strong predictor of genetic vulnerability to smoking (Hartz et al., 2012). Yet, even controlling for the respondent ‘s age of smoking initiation does not change the result.^4^

A causal interpretation of these results would suggest that some aspects of childhood SES reduce the influence of genetic factors. However, correlation is not causation: SES is not randomly assigned, and may be correlated with unobserved characteristics of the individual and her family (Plomin, 1990, 2014). Reassuringly, we find no evidence of correlation between thePGS and the four measures of childhood SES background (see SI section A.2). The absence of correlation between genetic risk and childhood SES background suggests the observed G×E interaction does not reflect a spurious gene-environment correlation (rGE) or a non-linear effect of genetic risk *G* that results from correlation between *G* and *E*. Other interpretations are still possible. For example, if genetic risk *G* is correlated with an unobserved environment *E*^*\**^ (e.g., “family social and cultural norms”) and *E*^*\**^ in turn is correlated with the observed environment *E* (e.g., “family well off”), then the effect of the observed environment *E* may not be causal,but rather proxy for the existence of an interaction between *G* and the unobserved environment *E*^*\**^. Since *E* is correlated with *E*^*\**^, our analyses still provide evidence for a true causal G×E interaction between max CPD and some environment that is correlated with childhood SES. However, in this example, improving the financial health of low-SES households with children may not improve later-life outcomes, since it might be family social and cultural norms, rather than financial resources, that matter.^5^

## 3 Discussion

We have provided empirical evidence that socioeconomic circumstances in childhood, or correlates of such childhood socioeconomic circumstances, substantially offset the genetic risk of heaviness of smoking. Therefore, policies targeting childhood circumstances have potential for reducing the genetic risk of peak smoking. Future work may wish to exploit natural experiments, providing exogenous variation in specific childhood socioeconomic circumstances, to establish what specific aspects of such circumstances causally protect against the genetic risk for peak smoking.

Understanding environmental exposures and critical periods of life that affect unhealthy behavior for different genetically at risk groups may lead to personalized approaches in which genetic information is used preventively, informing individuals to avoid environments that may harm them. From the perspective of private citizens, personalized genetically informed treatment is becoming a reality, as exemplified by the growth of the direct to consumer genetic testing industry, such as 23andMe, FitnessGenes, UBiome, DNAFit, Orig3n and Habit. From a public perspective, naturally we should beware of government policies targeting groups based on genetic information. But policymakers do not need to know the genetic makeup of individuals to develop policy (Belsky and Israel, 2014). For example, if social-science genetic studies, such as ours, find that smokers are solely genetically at-risk individuals, one might be limited to pharmacotherapy or to pharmacogenetics interventions (Hall et al., 2005; Nutt, 2007). But, if these are the genetically at-risk individuals who have experienced adverse (childhood) environments, prevention efforts targeting modifiable characteristics of such environments may reduce later-life dependence. Our results suggest modifiable characteristics of the childhood environment exist that protect against the genetic risk for peak smoking.

Our study further demonstrates that early socioeconomic circumstances are important and can have long-lasting effects. Disparities between SES groups in smoking, and more generally in health behaviors and health, may in part be the result of socioeconomic backgrounds exacerbating or moderating genetic risk.

## 4 Materials and methods

### 4.1 Data

The Health and Retirement Study (HRS) is a longitudinal household survey providing rich data on about 26,000 individuals, representative of the U.S. population over the age of 50. Up to12 waves, over 24 years, of data per respondent are available. We use the publicly available HRS core survey and linked genetic data for the years 2006, 2008, and 2010. HRS core surveys are conducted biennially using a combination of face-to-face and telephone interviewing. A random subset of the about 26,000 total participants was selected to participate in enhanced face-to-face interviews and saliva specimen collection (for DNA) in 2006, 2008, 2010, and 2012. Genotyping was conducted from 2011-2014, using the Illumina Human Omni-2.5 Quad beadchip (HumanOmni2.5-4v1 array), with coverage of approximately 2.5 million SNPs. To increase SNPs coverage, we used the best-guess genotypic data which was imputed using approximately 21 million DNA variants from the 1000 Genomes Project, phase I (The 1000 Genomes Project Consortium et al., 2012).

### 4.2 Childhood SES

A set of retrospective questions about an individual ‘s household circumstances during childhood were collected in the HRS. We dichotomized all SES variables, assigning a value of 1 for high childhood SES and 0 otherwise. The four variables we construct are:

- *Family Well Off*: High SES indicates respondents who reported that their family was “pretty well off financially” or “average” from birth to age 16. Low SES indicates respondents who reported that their family was “poor.”
- *Never Move or Ask for Help*: The HRS asks separate questions about whether a respondent ‘s family ever had to move residences or ask relatives for help due to financial reasons.Both measures capture extraordinary financial hardship, and therefore we combine them into a single variable to improve overall frequency.^6^ High SES indicates respondents whose family never had to move or ask relatives for help for financial reasons. Low SES indicates respondents whose families did either move or asked relatives for help.
- *Father Employed*: High SES indicates respondents whose father never experienced a significant unemployment spell of “several months or more.” Low SES indicates respondents whose father did experience a significant unemployment spell, or those whose fathers were dead or never lived with them.^7^
- *High SES Index*: We construct a simple index by summing all three measures, taking values {0, 1, 2, 3 }. We then divide the sample into low SES (index values of {0, 1 }) and high SES (index values of {2, 3 }). The latter dichotomous measure is the main SES index used in the analyses.

### 4.3 Polygenic Score

The polygenic score *PGS*_*i*_ is constructed using the software Plink (Purcell et al., 2007) as the weighted sum of *g*_*ij*_, the genotype of individual *i* for SNP *j* = 1, …, *J*.

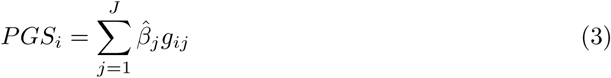

where 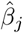 are the GWAS-estimated additive effect sizes of the alleles of SNP *j*, coded as having 0, 1, or 2 instances of the allele which is positively correlated with the phenotype (Dudbridge, 2013). No clumping or pruning was performed. We calculate the simple weighted sum of all SNPs present in the imputed HRS data set for which we have GWAS estimated additive effectsizes 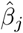 and that have a GWAS *P*-value of less than 10^−4^ (chosen to focus on SNPs with astrong association).

## Aknowledgments

Research reported in this publication was supported by the National Institute on Aging of the National Institutes of Health under Award Numbers K02AG042452, R01AG037398, and RF1AG055654. The content is solely the responsibility of the authors and does not necessarily represent the official views of the National Institutes of Health. Titus Galama is grateful to the School of Economics of Erasmus University Rotterdam for a Visiting Professorship in the Economics of Human Capital. We thank Marco Angrisani, Jonathan Beauchamp, Dan Benjamin, Donna Gilleskie, Donald Kenkel, and Hans van Kippersluis for helpful comments and discussions.

## A Supporting Information

### A.1 Analysis Sample

Table 2 provides summary statistics for the sample of the 3,280 respondents with non-missing information for all of the variables used in the regression of equation 1.

**Table 2:** Summary Statistics

| Variable | Mean | Std. Dev. | N |
| --- | --- | --- | --- |
| Outcomes: |  |  |  |
| Max CPD | 25.05 | 17.90 | 3,280 |
| Low SES Index & Low PGS | 26.76 | 18.02 | 374 |
| Low SES Index & High PGS | 30.66 | 19.66 | 377 |
| High SES Index & Low PGS | 23.15 | 16.86 | 1248 |
| High SES Index & High PGS | 24.74 | 17.95 | 1281 |
| Genetic Measures: |  |  |  |
| PGS | 0.00 | 0.98 | 3,280 |
| SES Measures: |  |  |  |
| Fam. Well Off | 0.73 | 0.44 | 3,280 |
| Father Emp. | 0.75 | 0.43 | 3,280 |
| Nev. Mov/Ask | 0.75 | 0.43 | 3,280 |
| SES index | 0.77 | 0.42 | 3,280 |
| Demographics: |  |  |  |
| Male | 0.53 | 0.50 | 3,280 |
| Birth year | 1939.46 | 8.82 | 3,280 |

“Low PGS” refers to negative PGS (below the mean); “High PGS” refers to positive PGS (above the mean).

Table 3 shows the polychoric correlations^8^ between the four measures of childhood SES, the outcome variable (max CPD), and the PGS (standard errors in parentheses).

**Table 3:** Correlations Between SES Measures and the PGS

|  | Max<br>CPD | PGS | Fam.<br>Well Off | Nev. Mov/<br>Help | Father<br>Emp. | SES<br>Index |
| --- | --- | --- | --- | --- | --- | --- |
| Max CPD | 1.000<br>(0.000) |  |  |  |  |  |
| PGS | 0.066<br>(0.016) | 1.000<br>(0.000) |  |  |  |  |
| Fam. Well Off | -0.132<br>(0.022) | 0.028<br>(0.024) | 1.00<br>(0.000) |  |  |  |
| Nev. Mov/Help | -0.119<br>(0.023) | 0.022<br>(0.024) | 0.538<br>(0.024) | 1.000<br>(0.000) |  |  |
| Father Emp. | -0.124<br>(0.023) | 0.013<br>(0.024) | 0.507<br>(0.025) | 0.574<br>(0.026) | 1.000<br>(0.000) |  |
| SES Index | -0.148<br>(0.023) | 0.015<br>(0.025) | 0.857<br>(0.012) | 0.896<br>(0.009) | 0.879<br>(0.011) | 1.000<br>(0.000) |

Pairwise polychoric correlations among the main variables of interest. Standard error in parenthesis.

### A.2 Correlation Between Measures of SES and the PGS

The correlation between the four measures of childhood SES and the PGS is small and indistinguishable from zero (see SI Table 3), and as SI Figure 2 shows, the distributions of the PGS for max CPD are remarkably similar for high and low childhood SES backgrounds. Quantitative analyses also provide no evidence of correlation. KolmogorovSmirnov tests for differences between the high and low SES distributions of the PGS for each of the four SES measures fails to reject equivalence with *P*-value=0.711 for the SES index, *P*-value=0.274 for family well off, *P*-value=0.327 for never move or ask for financial help, and *P*-value=0.730 for father employed (*P*-values shown in SI Figure 2). Further, SI Figure 3 shows the coefficients of quantile regressions testing the difference between high and low SES at every 5^*th*^ percentile of the distribution of PGS scores. The graphs show the 95^*th*^ percent confidence interval (without controlling for multiple hypothesis testing), indicating virtually no statistically significant difference at any percentile, for each of the four measures of childhood SES.

**Figure 2:**
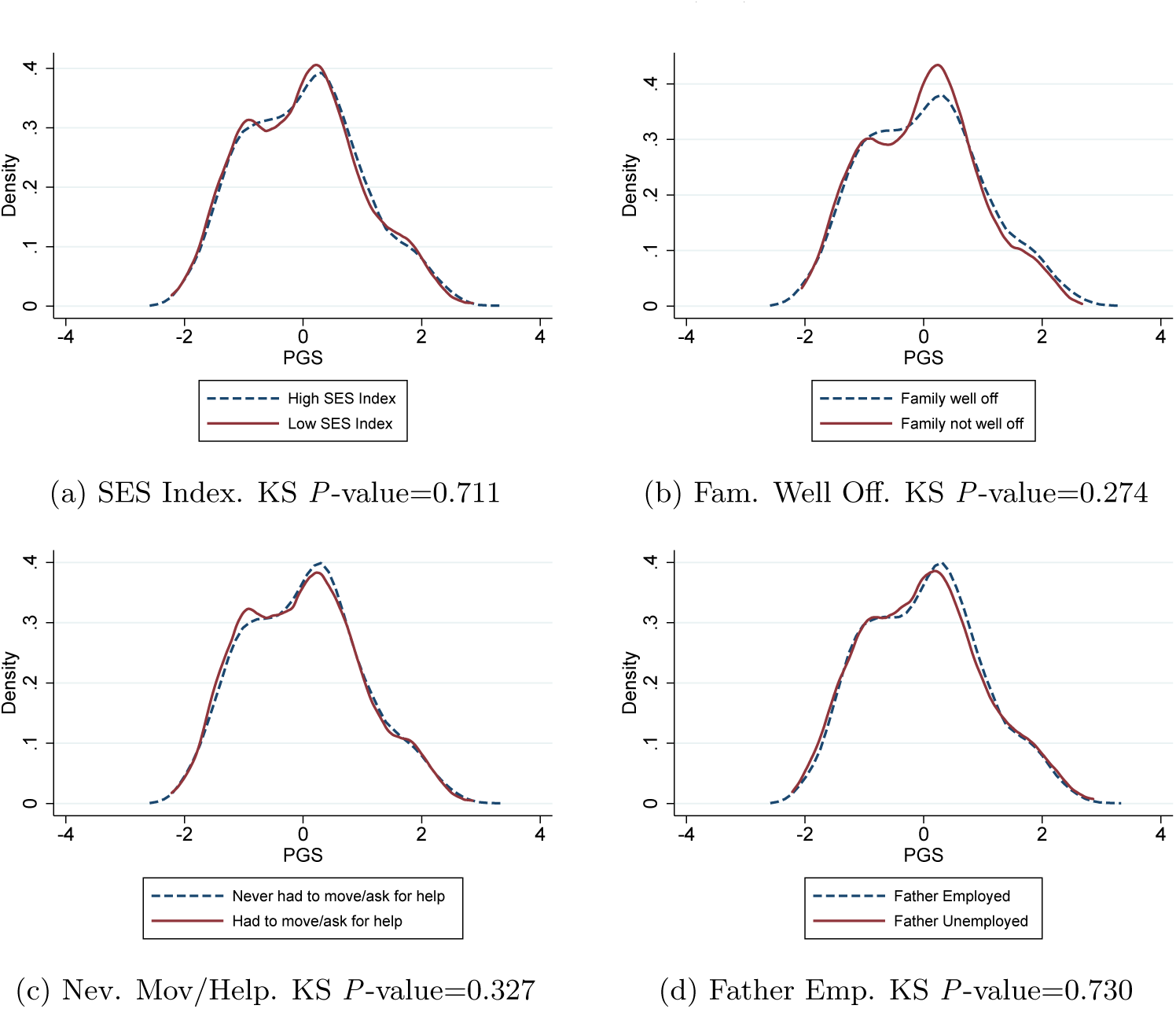
Epanechnikov kernel density distributions of the PGS for high and low SES and for different SES measures. Kolmogorov-Smirnov test for equality of distributions (KS) *P*-value shown.

**Figure 3:**
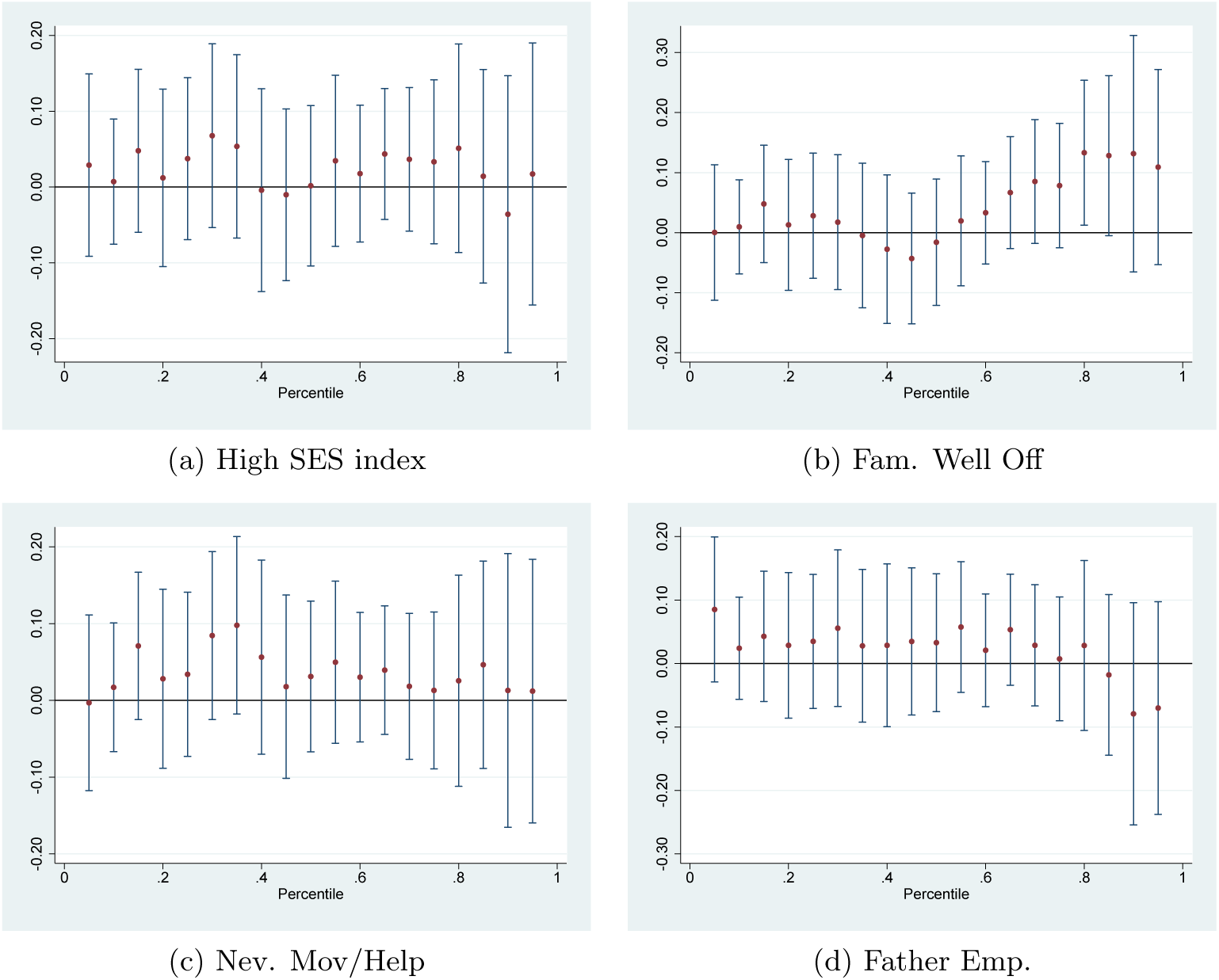
Differences in quantiles of the PGS distribution between high and low SES. This figure plots the coefficients from quantile regressions of the PGS over different SES measures. A separate regression was ran for every 5^*th*^ quantile of the PGS distribution, and for every measure of childhood SES. 95% confidence intervals shown.

Last, correlation between the PGS and SES is also very weak in the full sample where we do not restrict to smokers (see SI Figure 5). Therefore the lack of gene-SES correlation is not a result of our sample selection.

### A.3.Additional Analysis

Using the same procedure as was used in constructing Figure (1) but for the other three measures of SES, Figure (4) demonstrates G×SES interaction also in the “family well off” measure of SES as evidenced by the difference in slopes. On the other hand, for the “moving/asking for help” measure of childhood SES there are G and SES effects, but little evidence for G×SES interaction. For the “father employed” measure of childhood SES, a more complex pattern emerges with the sign of the G×E interaction reversing for those with higher genetic risk. As these analyses ofthe raw data demonstrate, linear regressions of the standard form 1 may fail to uncover G×SES interaction while present, and hide complex interaction effects.

**Figure 4:**
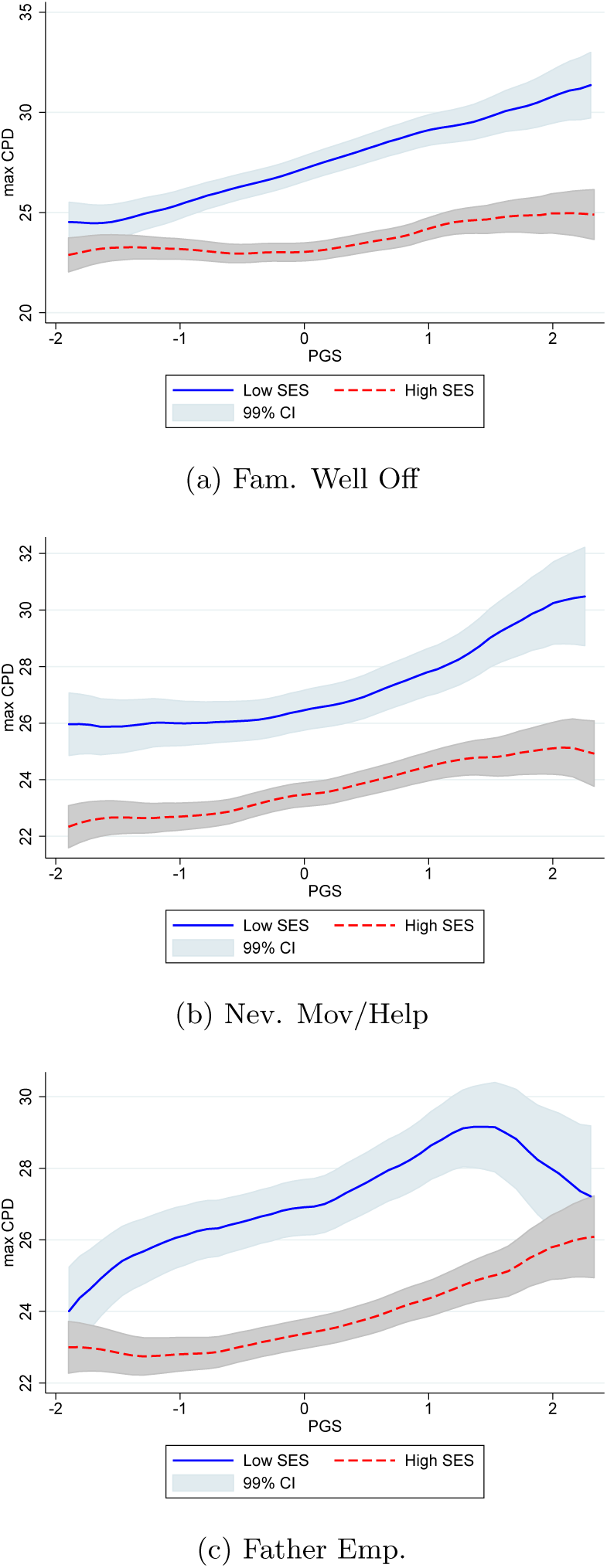
Smoking and polygenic score by different measures of SES.

**Figure 5:**
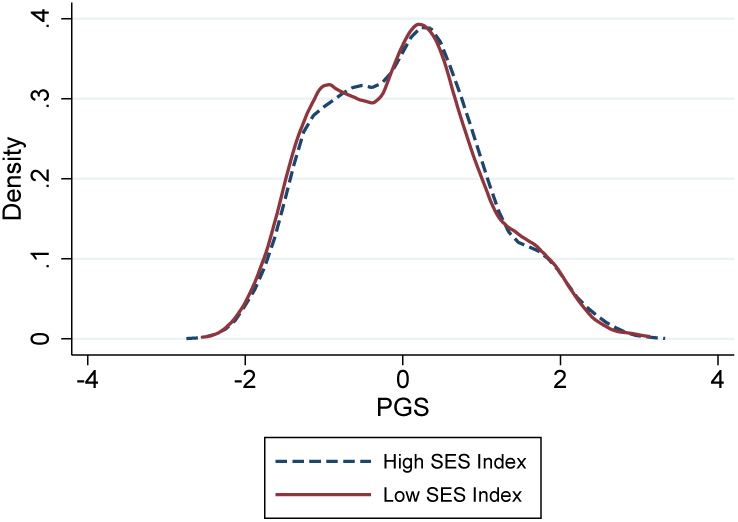
Distribution of Smoking PGS by SES, for the whole sample, i.e. not restricted to smokers. Kolmogorov Smirnov test for equality of distributions *P*-value=0.085.

Table 4 shows the results of estimating equation 1 separately by gender. Results are stronger for women, showing a slightly higher estimated coefficient for the direct SES and genetic effects and for the interaction term, but none of these are statistically significantly different from men.

**Table 4:** Interplay between PGS and childhood SES in max CPD by sex

| SES Measure: | Max cigarettes per day |  |  |  |
| --- | --- | --- | --- | --- |
|  | (1) | (2) | (3) | (4) |
|  | Fam. Well Off | Nev. Move/Ask | Father Emp. | SES Index |
| **Panel A: Male Sample** |  |  |  |  |
| High SES | -2.469**<br>(1.095) | -2.730**<br>(1.165) | -2.605**<br>(1.148) | -3.362***<br>(1.177) |
| PGS | 2.420***<br>(0.931) | 1.991**<br>(0.987) | 0.344<br>(0.986) | 1.698*<br>(1.017) |
| PGS × SES | -2.100*<br>(1.099) | -1.385<br>(1.151) | 0.877<br>(1.153) | -1.002<br>(1.166) |
| Controls | ✓ | ✓ | ✓ | ✓ |
| Obs. | 1,748 | 1,748 | 1,748 | 1,748 |
| $R^2$ | 0.206 | 0.206 | 0.205 | 0.207 |
| **Panel B: Female Sample** |  |  |  |  |
| High SES | -4.335***<br>(1.077) | -4.932***<br>(1.079) | -4.524***<br>(1.060) | -5.752***<br>(1.123) |
| PGS | 1.713*<br>(1.029) | 2.436**<br>(0.961) | 2.780***<br>(0.942) | 2.727***<br>(1.007) |
| PGS × SES | -1.005<br>(1.153) | -2.090*<br>(1.098) | -2.629**<br>(1.077) | -2.342**<br>(1.127) |
| Controls | ✓ | ✓ | ✓ | ✓ |
| Obs. | 1,532 | 1,532 | 1,532 | 1,532 |
| $R^2$ | 0.244 | 0.249 | 0.249 | 0.252 |

Coefficients estimated from OLS regression, separate by gender and by SES measure, of max CPD on SES, the PGS, the interaction between PGS and SES, and our baseline controls $Z_i$ (except for terms containing male indicators). Robust standard error in parenthesis. \*\*\* $P$ -value $< 0.01$ , \*\* $P$ -value $< 0.05$ , \* $P$ -value $< 0.10$

Table 5 reports the results of estimating equation 1 using different measures of childhood SES. The coefficient on the interaction term, *ρ*, is consistently negative, and statistically significant for three out of the four childhood SES measures, Father ‘s Employment being the only coefficient indistinguishable from zero. Therefore, our results do not hinge on the specific measure of childhood SES that can be considered.

**Table 5:** Interplay between PGS and various measures of childhood SES in max CPD

| SES Measure: | Max cigarettes per day |  |  |  |
| --- | --- | --- | --- | --- |
|  | (1) | (2) | (3) | (4) |
|  | SES Index | Fam. Well Off | Nev. Move/Ask | Father Emp. |
| SES | -4.460***<br>(0.771) | -3.412***<br>(0.737) | -3.720***<br>(0.753) | -3.669***<br>(0.746) |
| PGS | 2.477***<br>(0.685) | 2.367***<br>(0.655) | 2.237***<br>(0.657) | 1.672**<br>(0.654) |
| PGS × SES | -1.866**<br>(0.781) | -1.768**<br>(0.758) | -1.593**<br>(0.761) | -0.846<br>(0.760) |
| Controls | ✓ | ✓ | ✓ | ✓ |
| Obs. | 3,280 | 3,280 | 3,280 | 3,280 |
| $R^2$ | 0.197 | 0.194 | 0.194 | 0.193 |

Coefficients estimated from OLS regression of max CPD on SES Index, the PGS, the interaction between PGS and SES Index, and controls $Z_i$ . Robust standard error in parenthesis. \*\*\* $P$ -value $< 0.01$ , \*\* $P$ -value $< 0.05$ , \* $P$ -value $< 0.10$ .

Finally, Keller (2014) suggests that, in order to properly estimate G × E interaction, it is important to fully interact the genetic measure (PGS) and the environmental measure (childhood SES) with the full set of controls. The results of this procedure are shown in Table 6. The point estimate of the G × E interaction remains remarkably stable, even when fully interacted, albeit less precisely estimated. However, we may be over-fitting the data. The set of controls *Z*_*i*_includes 49 indicators for each year of birth (yob), each interacted with gender, the first ten principal components of the full matrix of SNP data, and a set of ten indicators for region of birth, amounting to a total of more than 400 control variables. Interacting all of these with both the PGS and the SES Index leads to more than 1200 controls. Not surprisingly, using this many controls in a sample of 3,280 individuals, the interaction effect is somewhat less stable.

**Table 6:** Interplay between PGS and high SES index in max CPD, fully interacted model

|  | Max cigarettes per day |  |  |  |
| --- | --- | --- | --- | --- |
|  | (1) | (2) | (3) | (4) |
|  | Baseline Model | Interacted Model |  |  |
| High SES index | -4.465***<br>(0.771) | -4.424***<br>(0.777) | 5.498<br>(15.025) | 2.930<br>(15.698) |
| PGS | 2.477***<br>(0.685) | 3.477<br>(6.386) | 2.229***<br>(0.707) | 2.385<br>(6.636) |
| PGS × SES | -1.866**<br>(0.781) | -1.615**<br>(0.802) | -1.533**<br>(0.803) | -1.224<br>(0.827) |
| Controls: |  |  |  |  |
| Demographic | ✓ | ✓ | ✓ | ✓ |
| Demo. × PGS | | ✓ | | ✓ |
| Demo. × SES | | | ✓ | ✓ |
| Obs. | 3,280 | 3,280 | 3,280 | 3,280 |
| $R^2$ | 0.197 | 0.217 | 0.216 | 0.235 |

Coefficients estimated from OLS regression of max CPD on SES Index, the PGS, the interaction between PGS and SES Index, and various controls. *Demographic* refers to the baseline control vector $Z_i$ ; *Demo. × PGS* are interactions between the baseline control vector $Z_i$ and the PGS. *Demo. × SES* are interactions between the baseline control vector $Z_i$ and the SES. Robust standard errors shown in parenthesis. \*\*\* $P$ -value $< 0.01$ , \*\* $P$ -value $< 0.05$ , \* $P$ -value $< 0.10$

To avoid over-fitting, we next use a more parsimonious model which includes a fourth-degree polynomial in year of birth instead of a separate indicator for each year of birth. The resulting control vector 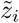 thus includes the first ten principal components of the full matrix of SNP data, a male indicator, yob, yob2, yob3, yob4, a set of indicators for the 11 regions of birth,interactions between the male indicator and the fourth-degree polynomial in yob, between the male indicator and region of birth indicators, and between the region of birth indicators and the fourth-degree polynomial in yob. As shown in columns (1) and (2) of Table 7, the estimated G× E coefficient is very similar regardless of whether we include the baseline control vector *Z*_*i*_ or the more parsimonious control vector 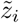. Interacting 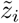 with both the PGS and the SES Index,the estimated G × E coefficient remains both statistically significant and remarkably similar in size to our baseline specification. Thus, our results are robust to fully interacting all controls with both the PGS and the SES indicator.

**Table 7:** Interplay between PGS and high SES index in max CPD, parsimonious controls

|  | Max cigarettes per day |  |  |  |  |
| --- | --- | --- | --- | --- | --- |
|  | (1) | (2) | (3) | (4) | (5) |
|  | Baseline Model | Parsimonious Model | Interacted Model |  |  |
| High SES | -4.465***<br>(0.771) | -4.211***<br>(0.727) | -4.138***<br>(0.732) | -7.662<br>(4.739) | -7.271<br>(4.780) |
| PGS | 2.477***<br>(0.685) | 2.539***<br>(0.643) | 0.738<br>(2.007) | 2.540***<br>(0.649) | 0.918<br>(2.019) |
| PGS × SES | -1.866**<br>(0.781) | -1.743**<br>(0.731) | -1.594**<br>(0.746) | -1.737**<br>(0.738) | -1.591**<br>(0.752) |
| Controls: |  |  |  |  |  |
| $Z_i$ | ✓ | | | | |
| $\tilde{z}_i$ | | ✓ | ✓ | ✓ | ✓ |
| $\tilde{z}_i \times$ PGS | | | ✓ | | ✓ |
| $\tilde{z}_i \times$ SES | | | | ✓ | ✓ |
| Obs. | 3280 | 3280 | 3280 | 3280 | 3280 |
| $R^2$ | 0.197 | 0.099 | 0.104 | 0.104 | 0.109 |

Coefficients estimated from OLS regression of max CPD on SES Index, the PGS, the interaction between PGS and SES Index, and various controls. $Z_i$ refers to the baseline control vector $Z_i$ , which contains birth-year fixed effects; $\tilde{z}_i$ refers to the parsimonious control vector $\tilde{z}_i$ , which contains a fourth order polynomial in birth-year instead of birth-year fixed effects; $\tilde{z}_i \times$ PGS are interactions between the parsimonious control vector $\tilde{z}_i$ and the PGS. $\tilde{z}_i \times$ SES are interactions between the parsimonious control vector $\tilde{z}_i$ and the SES. Robust standard errors shown in parenthesis. \*\*\* $P$ -value $< 0.01$ , \*\* $P$ -value $< 0.05$ , \* $P$ -value $< 0.10$

## Footnotes

1 They find the association between genetic risk and cigarettes smoked per day to be larger for those experiencing traumatic events and smaller for those who live in neighborhoods with greater social cohesion, using a small (∼1,500) sample of African Americans. Other studies evaluate the existence of G× E interplay for another health outcome, obesity. These find that unfavorable SES conditions amplify the genetic influence of 32 obesity-related genetic variants on body mass index (BMI), especially for recent birth cohorts (Liu and Guo, 2015); that years of schooling protect for genetic risk in type-2 diabetes (Liu et al., 2015); and that the social environment moderates the genetic influence of a variant near the FTO gene on the development of obesity in children (Foraita et al., 2015).

2 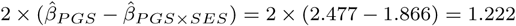. Additional analyses, reported in SI Table 4, shows little evidence for differences by gender.

3 This result is obtained from a regression of max CPD on an indicator equal to one for those reporting a degree from a four-year college, a masters, or a professional degree, and using the baseline control vector *Z*_*i*_.

4 Additional analyses, reported in SI Table 5, show that the coefficient on the interaction term, *ρ*, is consistentlynegative, and statistically significant for three out of the four childhood SES measures.

5 A more significant threat to a causal interpretation of our result occurs in a situation where genetic risk *G* is correlated with an unobserved environment *E*\*, and the true relation contains interactions between *E* and *E*\*. Consider the situation where the true data generating process is the following: 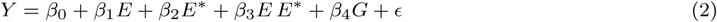 We observe *G*, but *E*\* = *αG* + *v*. The data generating process can be expressed as: 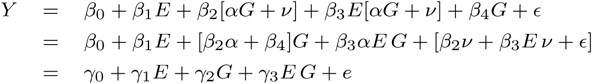 where the expectation of *γ*_3_ equals *β*_3_*α*, which is the strength of the *E* and *E*\* interaction and not the *E* and *G* interaction.

6 About 18 percent of respondent families had to move, and about 13 percent had to ask for help. When combined, about 25 percent had to take at least one of these actions.

7 This variable incorporates information on family structure since it takes the value 0 if the child is raisedwithout a father.

8 Polychoric correlations are used to estimate the correlation between continuous and dichotomous variables, such as our measures of SES. The STATA command *polychoric* has been used.

